# A neurocomputational model of the basal ganglia for the analysis of motor deficits after dopaminergic loss in Parkinson’s disease

**DOI:** 10.1101/2021.09.24.461656

**Authors:** Ilaria Gigi, Rosa Senatore, Angelo Marcelli

## Abstract

The basal ganglia (BG) is part of a basic feedback circuit, regulating cortical function, such as voluntary movement control, via their influence on thalamocortical projections. BG disorders, namely Parkinson’s disease (PD), characterized by the loss of neurons in the substantia nigra, involve the progressive loss of motor functions. At the present, PD is incurable. Converging evidence suggests the onset of PD-specific pathology prior to the appearance of classical motor signs. This latent phase of neurodegeneration in PD is of particular relevance in developing more effective therapies by intervening at the earliest stages of disease. Therefore, a key challenge in PD research is to identify and validate markers for the preclinical and prodromal stage of the illness.

We propose a mechanistic neurocomputational model of the BG at mesoscopic scale to investigate the behavior of the simulated neural system after several degrees of lesion of the substantia nigra, with the aim of possibly evaluating which is the smallest lesion compromising motor learning. In other words, we developed a working framework for the analysis of theoretical early-stage PD. While simulations in healthy conditions confirm the key role of dopamine in learning, in pathological conditions networks predict that there may exist abnormalities of motor learning process for physiological alterations in the BG which do not yet involve the presence of symptoms typical of the clinical diagnosis. Our model may account for the discovery of markers for an early diagnosis of the disease and give directions for developing novel noninvasive support systems.

## 1. Introduction

Even simple motor tasks involve a complex series of movements. Besides, skilled performance requires the coordination of the successive movements into a smooth and structured sequence so that they appear coordinated with each other. Deciphering the mechanisms by which brain structures generate this intricate motor output is a central challenge in neuroscience. Based on a wealth of data, the basal ganglia (BG) is thought to play a principal role in this path. The BG is subcortical nuclei part of several anatomical and functional loops, involving the cerebral cortex and the thalamus. It is involved in, among others, voluntary movements control, procedural learning, decision making, cognition and emotion. But the primary function is that of controlling and regulating the activities of the motor and premotor cortical areas for executing smooth movements after learning skilled responses to cortical inputs. Indeed, current theories of biological motor learning claim that early learning is controlled by dopamine-modulated plasticity in the BG that trains parallel cortical pathways through unsupervised plasticity at corticostriatal synapses. When systems-level interactions of multiple brain regions are involved, computational investigations provide a valuable complement to experimental brain research.

Modeling approaches could be applied to develop testable “computational biomarkers” to support diagnostic, prognostic, or treatment efforts, particularly. Considering that these brain structures are subcortical, their activity can be studied experimentally with an abstract level, through electrodes implanted in the depths of the brain and capture the LFP of groups of neurons; in contrast, cortical activity currently is more practically investigated and stimulated through intracortical microelectrode arrays (MEAs). Thus, this is a first reason to necessarily find other means, behavioral, clinical, noninvasive, to make predictions and research BG nuclei activity.

In particular, BG dysfunction is associated with Parkinson’s disease (PD), currently incurable and characterized by the loss of neurons in a region of the BG known as substantia nigra (SN). There is no accepted definitive biomarker of PD. An urgent need exists to develop early diagnostic biomarkers for two reasons: (1) to intervene at the onset of disease and (2) to monitor the progress of therapeutic interventions that may slow or stop the course of the disease.

We propose a biologically inspired mechanistic neural network model of the BG and learning rule in motor domain, incorporating known biological constraints. The goal is to generate a model that mimics information processing in the brain to simulate the performance of a certain motor task at some level of abstraction. Layers of interconnected neurons form brain networks, and the neurons are supposed to adjust their synapses to accommodate the appropriateness of the output to the task in a learning mechanism based on environmental feedback. Learning is at two different synapses: cortico-cortical and corticostriatal, and they are modeled differently. Exploration of the neural interactions could gain some insights about the functional dynamics of information processing within the simulated brain areas, in normal as well as in diseased brains, providing also some guidelines for developing more efficacious therapies for diseases in which these areas are involved.

A field of AI research deals with proving the existence of deteriorated behavior in handwriting as the expression of a BG lesion of small entity, in the early stage of the disease. In this light, our neurocomputational model enables to investigate the behavior of the simulated neural system after several degrees of lesion, to evaluate which is the smallest lesion compromising motor learning. Such a lesion would represent a marker of the disease, existing before the onset of evident symptoms, which occur once the SN is severely damaged. We additionally show that our model produces experimental and clinical findings that relate Parkinson’s disease, especially with respect to handwriting alterations caused by the disease.

## 2. Response selection in motor domain

The BG performs its action selection (choosing motor responses) function over a wide range of frontal cortical areas, by virtue of a sequence of parallel loops of connectivity; each region of the cortex has a corresponding BG circuit, for gating proper initiation of movement in that area. The motor circuit has been examined in experimental studies and has been implicated in a wide range of motor behaviors, particularly in the higher-level aspects of movement, such as the preparation of movement or the control of kinematic parameters. Basically, the BG integrates information from different brain structures into an appropriate motor response, e.g: in the reaction to a sensory stimuli or also in everydays movement and the fine control of all our movements, walking, writing and so on.

In the motor loop, the associated striatum (putamen) receives input primarily from motor cortex (MC), namely, supplementary motor area (SMA), cingulate motor area (CMA), premotor cortex (PMC), and primary motor (M1), and also from parietal cortex, mostly primary somatosensory cortex (S1). Then, the input signals come from a large region of the frontal-parietal areas, and are everything needed to properly contextualize motor actions. A high-level diagram of the motor loop is reported in Figure 1a. The inputs converging into the BG are there processed and reissued, through the thalamus, specifically the ventrolateral (VL) nuclei, to the cortex. The thalamus projects principally to frontal lobe areas (prefrontal, premotor and supplementary motor areas) which are concerned with motor planning. The BG serve as a gate on the thalamocortical circuit, in that it modulates, through direct and indirect pathway, the execution or inhibition of a motor command by acting on the thalamus, but the contents of the information that go through the gate (e.g., the specifics of the motor action plan, like the force or size of a movement) depend on the thalamocortical system. This fairly complex circuit is motivated by computational considerations and a lot of detailed anatomical, physiological, and pharmacological data, and it has been overwhelmingly supported by empirical evidence, in humans and other species, for example [1]. From the MC, ultimately the BG affects the planning and execution of the movement by synapsing with the neurons of the corticospinal and corticobulbar tracts in the brainstem and spinal cord.

**Figure 1:**
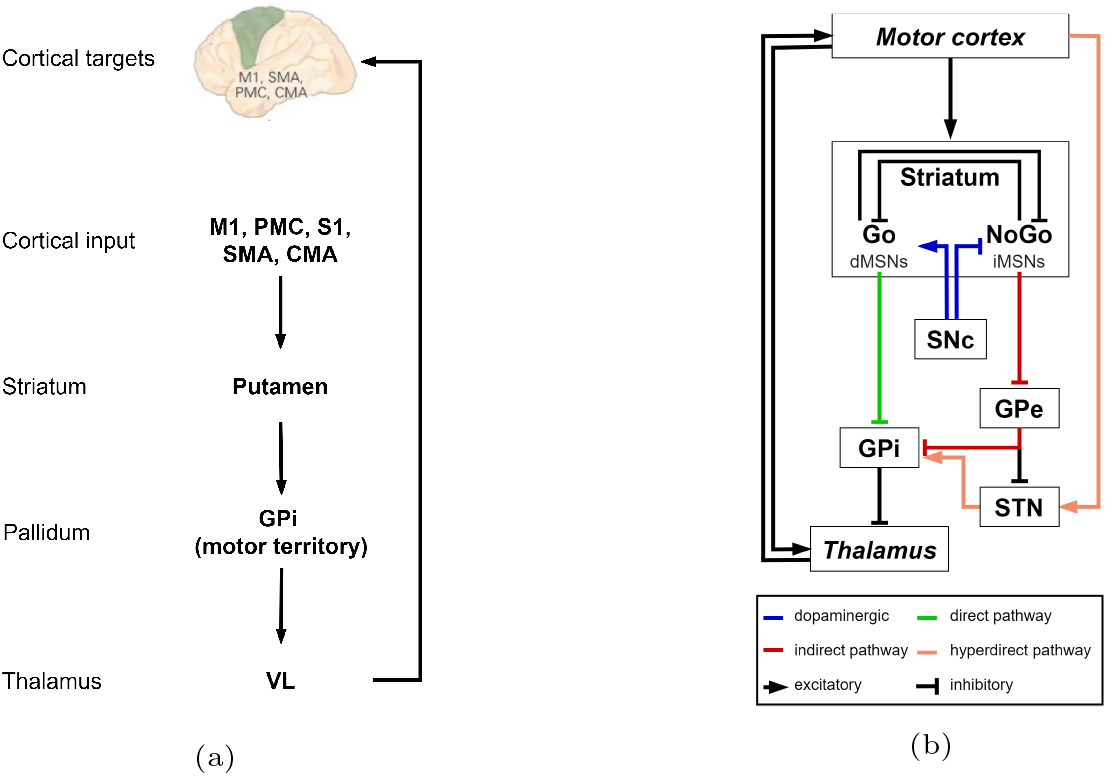
1a Basal ganglia functional loop: motor neurocircuit for body movement. 1b Functional description of the model. The corticostriato-thalamocortical loop, including the direct (DP), indirect (IP), and hyperdirect (HP) pathways of the BG is represented. The direct pathway-projecting medium spiny neurons (dMSNs) or Go neurons in the striatum project directly to the GPi, having the effect of disinhibiting the thalamus and executing a motor response represented in the cortex. The indirect pathway-projecting medium spiny neurons (iMSNs) or NoGo neurons have an opposite effect and suppress actions from getting executed. Dopamine from the SNc projects to the striatum, exerting excitation of Go cells via D1 receptors, and inhibition of NoGo via D2 receptors. The STN receives excitatory projections from the cortex in the hyperdirect pathway and excites GPi; GPe provides inhibitory effect on STN activity. Areas not belonging to BG are marked in italic.

Current theories of biological motor learning pose that early learning is controlled in the BG by dopamine-modulated plasticity, which trains parallel cortical pathways through unsupervised plasticity as a motor task becomes well-learned. In human trial-and-error learning tasks, phasic bursts and dips of dopamine (DA) have been proved to occur during positive and negative feedback, respectively, and these changes in extracellular levels of DA during feedback are thought to be critical for learning because they modify synaptic plasticity; in other words, DA acs as a *teaching signal*, leading to the learning of rewarding behaviors and discouraging unrewarding ones [2, 3, 4]. The circuitry that implements these functions is described next. Thirdly, it has been widely agreed that habit learning impairment in PD is linked to damaged DAergic neurons in the BG and DA deficiency [5, 6, 7].

## 3. Model description

The goal of the present study is to design a biologically plausible functional model of the BG, based on experimental findings and computational studies, to replicate voluntary motor initiation in a classical high-level motor learning task. To this end, this paper presents a computational model of the cortico-BG-thalamo-cortical loop, diagrammed in Figure 1b, in order to understand how a motor command is associated to a particular sensory state through a learning process and how neural elements interact to yield this emergent behavior at the level of neuronal layers; in other words, we propose a biophysically detailed mechanistic model at mesoscopic scale.

A commonly accepted hypothesis of BG interconnections with the cerebral cortex is the circuit formed by the premotor cortex (PMC), the primary motor cortex (M1), a cortical input signal provided by different motor areas, responsible for planning of complex voluntary movements and encoding states or categories, the thalamus, and the BG. Therefore, the model includes a BG-thalamocortical closed loop with the MC.

The BG system involves the following subregions: striatum (D1 and D2 receptors), subthalamic nucleus (STN), substantia nigra pars compacta (SNc), the internal globus pallidus (GPi) and the external globus pallidus (GPe).

The model incorporates the principal aspects of the BG anatomy and neurobiology, together with cellular and systems-level effects of DA. It is included the direct, the indirect and the hyperdirect pathways, striatal interneurons, DA-mediated reinforcement learning rule to learn cortico-striatal synapses, and hebbian-based learning mechanism for cortico-cortical synapses. Each brain region has been modeled as a layer within the network. Neurons in the striatum layer are topographically-organized in columns to represent a segregated circuit for multiple actions to be modulated in parallel. The topography is also maintained across the BG output nuclei (GPe and GPi) and in downstream targets (thalamus, PMC, M1), which is in agreement with empirical studies confirming the parallel model of BG function (e.g, [8]).

The core theoretical features of the functional role of the BG nuclei in motor habit learning adopted in the model are listed herein and explained in depth in the following sections:

- The cortex generates multiple competing candidate actions for a given sensory context, which are kept inhibited by tonic activity in the output structures of BG. The striatum is able to release the inhibition on one of this channels, such that the most rewarding one is chosen to be activated.
- Competitive dynamics between striatal cells in the direct and indirect pathways of the BG facilitate or suppress a given response in the cortex. The cells that detect conditions to facilitate a response provide a *Go* signal, whereas those that suppress responses provide a *NoGo* signal.
- DA has a key role in modulating both activity and plasticity in the BG. Phasic changes (bursts and dips) in DA during error feedback are critical for learning stimulus-reward-response associations and for allowing Go/NoGo representations to actually facilitate or suppress the execution of a command.
- Less DA (as in PD state) leads to less contrast enhancement and impairs the ability to resolve Go/NoGo association differences needed for discriminating between different responses.
- Hebbian learning enhances associated cortical representations in a learning process slower than the reinforcement-based one.
- The STN provides a *global NoGo* signal that suppresses all responses. By this account, cortico-subthalamic-pallidal pathways modulate the dynamics of action selection by regulating the threshold for executing a response. This function is adaptively adjusted by the degree of cortical activation; put another way, the STN prevents impulsive or premature responding.

### 3.1. BG network functionality

The BG circuitry incorporates the (dorsal) striatum, namely putamen, the globus pallidus (internal, GPi, and external segments, GPe), the subthalamic nucleus (STN) and the substantia nigra pars compacta (SNc). In the context of motor control, the BG facilitates the execution of appropriate actions and suppress all the other, inappropriate, ones; in other words, the BG modulate motor responses rather than encoding any detail of them, *releasing the brake* [3] on the motor command winning the competition (i.e: getting executed) and represented in the motor/premotor cortex. Thus, in our model the BG is not an SR module, but instead modulates the efficacy of responses being selected in the cortex. This model suggests that the BG does not initiate motor responses, rather facilitates or gates responses that are being considered in the PMC [3]. Looking at it another way, the BG modulate elementary actions, which are then pieced together by the nervous system to generate a complex motor behavior, that results from the concatenation of elementary movements. In this complex system, the BG acts by enhancing the most appropriate command at any given portion of the sequence of actions and at the right time.

The input structure of the BG is the striatum/putamen and receives inputs from two main sources. First, multiple cortical areas, including the MC, project to the BG. Second, there is a DAergic projection from the SNc. Two main projection pathways go from the striatum to the output segment of BG, the GPi, up to the thalamus and back to the cortex, which have opposing effects on the excitation/inhibition of the thalamus (i.e: execution/suppression of the action). The direct pathway facilitates the execution of responses, whereas the indirect pathway suppresses them. Neurons from the direct pathway project from the striatum through an inhibitory connection to GPi, which is tonically active and inhibiting the thalamus. Thus, the excitation of the direct pathway results in the inhibition of the activity of GPi, which in turn ceases to inhibit the thalamus; the thalamus is enabled to get excited from other excitatory projections and thalamocortical projections propagate the excitation to the cortex, enhancing the activity of the motor response (i.e: the brake is released) currently represented in the MC, so that it can be executed. Neurons in the indirect pathway inhibit the GPe, tonically active and inhibiting the GPi. Therefore, the excitation of the indirect pathway releases the tonic inhibition of GPe onto GPi, which in turn is more active and then further inhibits the thalamus, suppressing actions from getting executed, thereby having an opposite effect (see Figure 1b for a pictorial description of this circuitry).

The output structure of the BG, the GPi, is thought of as acting like the volume dial on a radio because its output determines whether a movement will be weak or strong. The mechanism described above accounts for the interpretation, from a functional point of view, of direct pathway neurons sending a Go signal to execute a response and indirect pathway neurons a NoGo signal to suppress all the others.

It is still controversial whether the direct and indirect pathways compete with each other or are functionally independent. That is, ultimately only one of Go or NoGo pathways activity predominates in the excitation/inhibition of the GPi/thalamus, thus amplifying or decreasing the force of movement. Experimental findings suggest that the segregation between two subregions of neurons is not distinct as once thought. Here, our computational model adopts the clear distinction between two subpopulations of neurons in the striatum, based on differences in biochemistry and efferent projections, to analyse the dynamical activity of the different BG nuclei and of the overall system, which might be unfeasible with detailed anatomical models.

Consistently with experimental findings, we implemented two synaptic mechanisms that can mediate learning: reward-related plasticity of cortico-striatal synapses and activity-dependent Hebbian plasticity of cortico-cortical synapses, so that early in learning the BG learns to facilitate most rewarding responses associated to stimuli, in a trial and error fashion, and later, once the associations are ingrained and responses are executed rather then acquired, there is less need for the selective facilitation by BG. Thus, Hebbian learning drives the training of the associations between the states encoded in cortical input neurons and motor responses represented in premotor cortical neurons, enabling automaticity. Indeed, later in learning the expression of the behavior becomes DA-independent.

Substantial evidence provide insight into understanding how the STN participates in decision making, both in motor control and cognitive processes. The STN is part of the hyperdirect pathway, so called because cortical projections towards the STN directly excite GPi, bypassing the striatum. The cortico-STN pathway has been shown to have a substantial effect on modulating when a response is executed. Recalling that the GPi is tonically active, then the activation of STN by cortex enhances GPi activity and its inhibition on the thalamus, therefore less likely to facilitate a response. Its effects are dynamic as response selection process evolves, and its efficacy depends on cortical excitatory projections. This connection is spread from the STN to the GPi, therefore the hyperdirect pathway represents a non-specific (with respect to response) excitatory process to cancel (all) inappropriate actions. Thus, increased cortical activity leads to dynamic adjustments in the timing to reach a response threshold for the selection, therefore influencing which response is ultimately facilitated. Indeed, Frank predicts that the STN may be essential to allow all information required to make decisions to be integrated before facilitating one, and thereby prevents premature responding [9]. In other words, the global NoGo signal provided by the STN changes to allow the model sufficient time to consider all the possibilities before selecting the most appropriate response.

To recap, given current input state and past experience, the model selects a response through the interaction of distinct parallel subloops, each modulating a response. Each subloop comprises a direct (Go) and an indirect (NoGo) pathway for a given response. The STN provides a global modulatory NoGo signal on all responses, initially suppressing, and, later, dynamically facilitating the most appropriate response allowing one Go signal.

### 3.2. DA in the BG: effects on activity and synaptic plasticity

Dopamine (DA) plays a crucial role in human and animal cognition, substantially influencing a variety of processes including reinforcement learning, motivation, and working memory. DA modulates the balance of the two pathways by exciting activity in Go neurons, through D1 receptors, while inhibiting NoGo activity, through D2 receptors, in line with consistent physiological evidence. SNc is thought to be tonically active and to exert a strong influence on striatal excitability. Given this, it is straightforward to assume that it may play a role in corticostriatal synaptic plasticity. Actually, in this regard, compelling evidence has shown that DA is the central player in the induction of plasticity at corticostriatal synapses on striatal neurons, in concert with other neurotransmitters participating in this drama.

DA firing patterns fluctuate between two different modes: phasic and tonic. The phasic signal is fast-timescale and spans milliseconds, whereas the tonic signal is slow-timescale and can span minutes or hours. The functions of tonic DA remain unclear.

Various studies suggest that phasic changes in DA plays a key role in synaptic plasticity and reinforcement learning, as are thought to occur during error feedback (e.g., [10, 3]). These DA changes have also been inferred to occur in humans receiving positive and negative feedback in cognitive tasks and cause the two subpopulations of striatal neurons to independently learn positive and negative reinforcement values of responses.

In the context of reinforcement learning and action selection, studies on PD patients have suggested that phasic bursts (and dips) facilitating learning from positive (and negative) feedback are effective because of the large contrast enhancement during the feedback. In summary, increases in DA result in increased contrast enhancement in the direct pathway and suppression of the indirect pathway. Phasic dips in DA have the opposite effect, releasing the indirect pathway from suppression. Therefore, the phasic changes in DA are critical for modulating Go/NoGo representations in the BG and ultimately facilitate or suppress the execution of motor commands. In other words, DA phasic levels correspond remarkably well with a non-specific reward-mediated trial-by-trial training signal in reinforcement learning models, as it allows synapses active on trials when a correct rewarded response was produced to be strengthened and synapses active on error trials to be weakened.

Thus, too much DA in the BG, as in case of L-dopa-based medications in PD, would lead to excessive activation of the cortex and bias the direct pathway. In contrast, DA antagonists bias the indirect pathway, because a lack of DA in the BG would lead to too little excitation of the cortex.

In our model, positive feedback (i.e: correct response) is supplied by a transient increase in simulated DA, which consequently enhances synaptically-driven activity in the direct (Go) pathway and reduces activation in the indirect (NoGo) pathway, driving Go learning. Conversely, phasic dips in simulated DA follow incorrect responses and result in the release of indirect (NoGo) inhibition from DA, so that the NoGo neurons can be excited from corticostriatal afferents, driving NoGo learning. Hence, DA release following rewards results in longterm potentiation (LTP) in the direct pathway and long-term depression (LTD) in the indirect pathway. Conversely, DA dips following the absence of a reward produces LTD in the direct pathway and LTP in the indirect pathway.

To summarize, striatal Go/NoGo associations are learned through phasic changes in simulated DA firing during positive and negative feedback, resulting in modulation of synaptic plasticity and supporting learning. Bursts of DA lead the model to learn to facilitate rewarding behaviors and sharpen representations in the Go pathway. Dips of DA allows the model to learn NoGo representations to the incorrect responses and to suppress disadvantageous behaviors. Thus, information about the values of choices represented in the cortex is encoded through the strength of the cortical synapses, whereby stronger synapses amplify the activity of cells. This process is model-free DA-based reinforcement learning.

### 3.3. Reinforcement learning in Parkinson’s disease

Studies on PD indicate that the disease is characterized by the death of DAergic cells projecting to the BG, and associated motor symptoms including tremor, rigidity, and slowness of movement. Moreover, PD patients are also affected by a variety of cognitive deficits, in particular, in procedural learning and decision making. Various authors [9, 10, 11] agree on the role of BG in selectively facilitate the execution of a single elementary motor command, while suppressing all competing others. A further step has been taken by the hypothesis that the BG participate in cognitive decision making similarly to the context of motor control, since the loops linking the BG with frontal systems are very similar and because of cognitive deficits observed in PD patients [3, 12, 13, 14]; for this reason, the role of action selection can be naturally extended to include higher-level cognitive decisions.

On the other hand, in agreement with Graybiel and Grafton [15], after more than a century of work on the motor functions of BG, in the last decades, researchers have focused on the functions in relations to learning habits and acquiring motor skills. Much evidence [15, 16, 17, 18, 19] has been collected and reviewed suggesting a major role of BG in learning behaviors by refining action selection according to an optimization process and shaping skills as a modulator of motor repertoires; and, this learning mechanism supported by striatal circuitry generalize to other domains, including cognitive skills. There is also evidence linking PD to degradation of motor learning [20, 2, 21, 22].

In short, while PD patients are still able to learn, some aspects of learning are degradated and faulty, but the details are not clear yet. Much research is investigating in the features of motor degradation affecting individuals with PD, to determine specific therapies and technologies to partially counteract these symptoms of the disease. Manipulations of DA levels in the striatum in animal models are shown to affect the activation of behavior, also in the sensorimotor context and the development of skilled motor responses [23, 24].

Given the hypothesized role of BG in decision making learning both in motor and cognitive context and that PD patients have severely depleted levels of DA in the BG, the functional role of the BG and the DAergic system in motor and cognitive processes represents an issue of particular interest.

### 3.4. Summary of the theoretical model

The features incorporated by our mechanistic model and implemented in a neural network model to test their potential role in motor learning are:

- The model includes the competing processes of the direct and indirect pathway and whether to gate (Go) or suppress (NoGo) a response is learned.
- The model includes the SNc so that the role of DA can be implemented and manipulated, with simulated D1R and D2R in the striatum.
- The implementation of the model includes four competing responses, but can be extended proportionally to include more alternatives.
- The model incorporates the STN and thus allows to explore its contribution in providing a global modulatory signal on facilitation and suppression of all responses.
- The model implements STN dynamic effects, as response selection process evolves, both within the trial and with the advance of learning.
- The model take into account DA bursts and dips, with an increase of SNc unit firing for correct responses and a decrease to zero SNc firing for incorrect responses.
- The model simulates a learning rule based on a reinforcement learning paradigm inspired by the learning algorithm of Leabra [25], which is a combination of error-driven and Hebbian learning.
- The model incorporates M1 itself to learn to favor a given response for a particular input stimulus, via Hebbian learning from the input layer.
- The model embodies the procedural motor learning hypothesis discussed in Section 2, allowing the BG to initially learn which response to gate via phasic changes in DA, ensuing from random cortical responses, and then this learning transfers to the cortex. In other words, BG and DA critically mediate the acquisition of a behavior, but they play a diminishing role in executing well-learned behaviors.
- The model includes a naive functional model of striatal interneurons for intrastriatal inhibition, and their synapses are constant.

## 4. Materials and methods

This section describes the computational features of our BG model inter-connected with the other cortical (S1, PMC, and M1) and subcortical (thalamus) areas, including the key learning mechanisms, connectivity and general simulation methods. The intention is to explain how the model achieves the functionality it does, the motor loop for learning and regulating the execution of a motor behavior, and give the reader a foundation for understanding the simulations discussed in the Results section. In particular, we follow a validated model of decision making [3, 9, 26] and modify it for motor domain. For the sake of notation simplicity, we will identify with a generic input layer all the cortical regions we have discussed in Section 2 and involved in the reverberating activation loops between BG and thalamus. Four parallel loops are implemented through both the striatum, GPi, GPe, thalamus, PMC, and M1 in order to support four responses and these are easily extensible to accommodate more.

### 4.1. General methods

The model is implemented within the Neural Simulation Technology Initiative (NEST) framework using an implementation of a basic point-neuron conductance model, the conductance-based leaky integrate-and-fire (IAF) neuron model, which provides a standard biophysically grounded model of individual neuron dynamics [27]. Specifically, all the simulations were performed using the simulator NEST 2.20.0 [28].

The adoption of NEST enables to drive an in-depth analysis, more extensively than other simulators, as it allows to set (and also measure) the firing rate or pattern of a population of neurons and to examine in details the behavior of both the externally-manipulated neurons and the intermediate ones in the network in terms of their neuronal activity (i.e: firing rate, synapse strength, or timing delay). In this framework, our idea is to simulate neuronal lesions to various degrees, acting on the firing rate of neurons rather than simulating only the presence/absence of a neuron.

There are simulated excitatory and inhibitory synaptic input channels. Synaptic connection weights have been trained using a reinforcement learning algorithm, further details in this section. In short, the learning algorithm involves two phases, allowing simulation of feedback effects. In the “minus phase” the network settles into activity states on the basis of input stimuli and its synaptic weights, ultimately choosing a response; in this sense, it represents a response phase. In the “plus phase” the network resettles in the same manner, with the only difference being a change in simulated DA: an increase for correct responses, and a dip for incorrect responses. So, it embodies a feedback phase to represent the reward process. Connection weights are then adjusted to learn on the difference between activity states in the minus and plus phases.

A schematic of the theoretical cortico-basal ganglia-thalamo-cortical (CBGTC) loop architecture is shown in Figure 1b; the above anatomical and functional considerations are synthesized in a spiking neural network model (Figure 2). The model learns to select one of four responses to different input stimuli. Direct and indirect pathways enable the model to learn conditions that are appropriate for gating as well as those for suppressing. Differing subtypes of neurons are organized within separate layers with names as shown; within layers, parallel reverberating activation loops for each action are represented in a column of units; such subloops independently modulate each response, allowing selective facilitation of one response with concurrent suppression of the others. Projections from the SNc to the striatum incorporate modulatory effects of DA. Simulated phasic bursts and dips in SNc firing ensue from correct and incorrect responses, respectively, and drive learning by preferentially activating the direct pathway after a correct response and the indirect pathway after an incorrect response. The model is trained on simulated version of a long-term motor skill training task, further details in Task description section.

**Figure 2:**
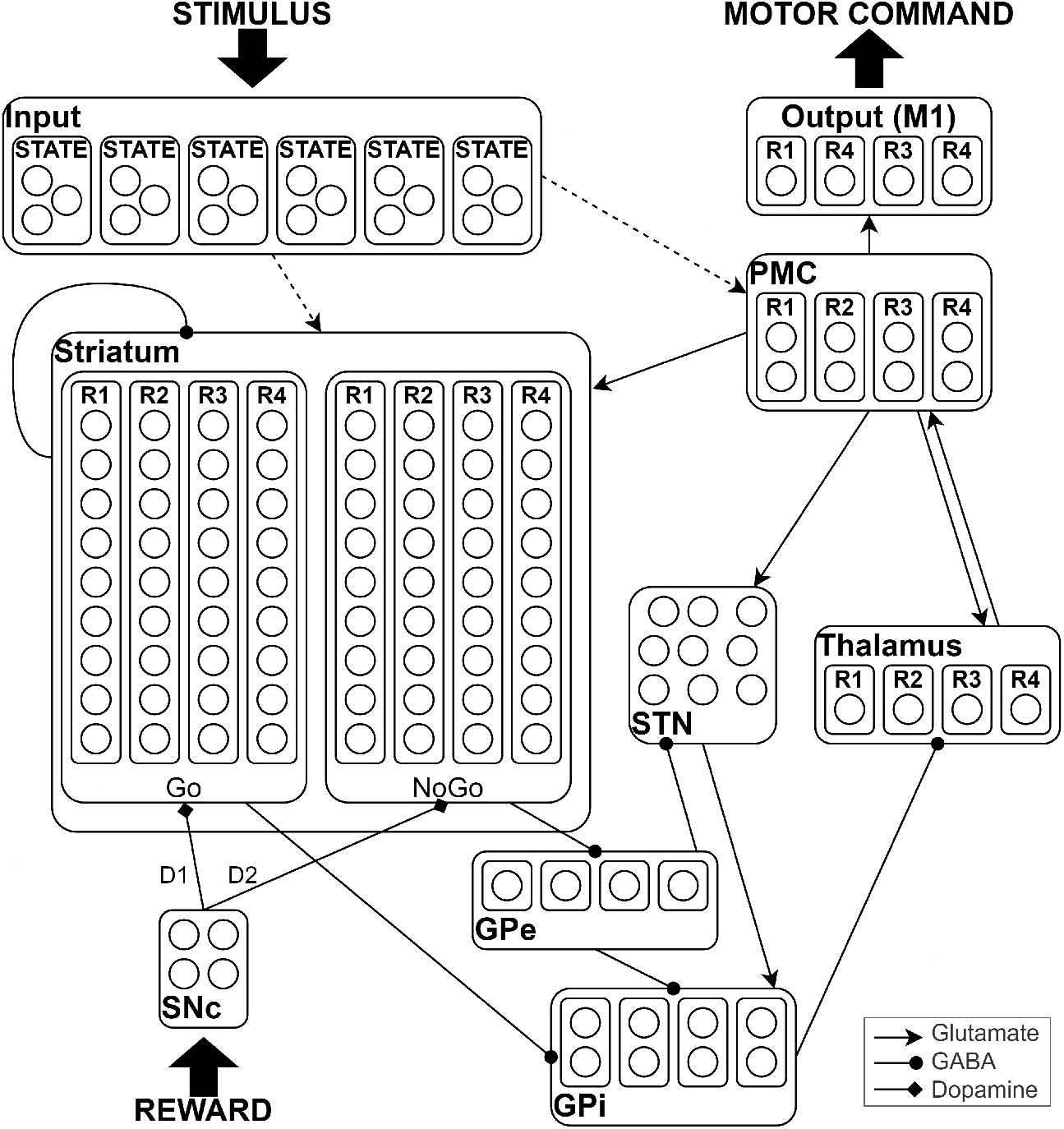
Network structure. Each rectangle represents one nucleus of the BG. Rectangles inside nuclei represent sub-groups or columns and circles stand for neurons. Dashed lines indicate plastic connections and solid lines stable connections not being learned. Arrow ends represent excitatory connections while dot ends inhibitory. The process of action selection is expressed by direct, indirect and hyperdirect pathways of the BG, with differential modulation of these pathways by DA in the SNc. The PMC selects a response via direct projections from the input (contrast enhancement in the direct (Go) pathway). Reward and cue generation were not explicitly implemented in the model but are shown above for completeness.

As we focus on the whole BG some simplifications are made with respect to computational details. Mainly, we do not model in detail the function of different interneurons and omit some known structural connections in the BG. Our reduced model of the BG consisted of 131 neurons. Number of neurons in each subpopulation and number of connections are provided in Table S1.

We are now describing the layers and their synaptic connections, referring to Figure 2, to guide the reader in a bottom-up approach through building up the network and the experimental setup. For the sake of clarity, we provide for each layer the description of its input/output synapses contextualized to their functionality, whereas we deal with exogenous sources in a separate subsection.

*Striatum*. In the neurophysiology, direct (Go) and indirect (NoGo) pathways start from two distinct populations of neurons in the striatum, expressing D1 and D2 receptors, respectively. In our neural network, the four leftmost columns of the striatum represent the direct pathways or Go (i.e: indexed 0-3), whereas the four rightmost columns represent the indirect pathways or NoGo (i.e: indexed 4-7). Each column is involved in the selection/inhibition of a particular response (R1–R4). The Go columns project to the corresponding columns in the GPi, whereas the NoGo columns project to the corresponding column in the GPe. Electrophysiological studies point out that interneurons are responsible for a tonic inhibitory activity in the striatum and are critical for shaping neuronal circuit activity in it, particularly for modulating the activity of medium-sized spiny neurons (MSNs). For this reason, we consider that to explicitly model interneurons as an additional functional module is essential. In our network, interneurons have been modeled as synaptically-weak inhibitory connections among striatal neurons.

*GPe*. GPe columns inhibit the associated columns in GPi, so that striatal Go and NoGo activity have opposing effects on the GPi. The inhibitory connection between cortex-striatum-GPe-GPi represents the indirect pathway.

*GPi*. Each column in the GPi tonically inhibits the associated column of the thalamus, which is reciprocally connected to the PMC. If Go activity is stronger than NoGo activity for a response, the corresponding column of the GPi will be suppressed, removing tonic inhibition of the corresponding thalamus unit, and eventually facilitating its execution in the MC, and it represents the direct pathway.

*STN*. In our model, the excitatory projections from the STN to the GPi convey cortical excitation and represent the hyperdirect pathway, which prevents premature choices suppressing movement through elevation of GPi activity. In addition, the other principal movement-suppressing pathway of the BG, the indirect pathway, interact within the STN with an inhibitory connection: all GPe units also inhibit STN neurons, in line with data showing that multiple GPe neurons converge on a single STN neuron. These synapses are stable, in line with studies showing that under normal conditions GPe-STN inhibition and cortical excitation is homeostatic, that is GPe-STN inhibition has been shown likely to limit cortical activation of STN units [29].

*SNc*. The BG nuclei receive the reward signal from the SNc through the nigrostriatal DAergic projections, which incorporate modulatory effects of DA. All the units of SNc excite Go columns (D1R) and inhibits NoGo columns (D2R).

*Cortex and thalamus*. In line with the hypothesis that the BG modulates the execution of “actions” (e.g., motor responses) being considered in different areas of the cortex, we consider a simple cortical hierarchy, consisting of a single input layer, conveying the ultimate aggregated information or encoding states, derived from different cortical contexts (sensory, motor programs, etc), and a two-level output hierarchy comprising PMC and M1. Two sets of synapses are plastic: (1) connections from the input cortex to the PMC (to learn category– response associations) and (2) connections from input cortex to the striatum (to learn stimulus–response associations). The (1) synapse is based on a Hebbian learning mechanism and (2) on DA-mediated reinforcement learning. The thalamocortical system models the reverberating activation from BG to cortex and is involved in stable synapses.

### 4.2. External inputs

The input stimulus is modulated through a simulated distinct activation of columns in the input layer, representing the condensed information of a contextualized environment state.

The reward signal generation mechanism is omitted, rather a tonic level of DA is simulated by setting the SNc units to be semi-active (0.5 spikes/ms) in the minus phase. Then, after the network selects a response, if the response is correct, a phasic increase in SNc firing occurs in the plus phase, with all SNc units set to have high firing rate (1.0 spikes/ms), causing a burst of DA, representing the reward. For an incorrect response, a phasic dip of DA occurs, with all SNc units set to zero activation (0 spikes/ms). In short, DA modulation has been structured as a rate-variable Poisson generator.

Throughout the network, neuronal populations receive external excitatory synaptic input according to Poisson’s distribution, to achieve realistic baseline firing rates and simulate both noise and background activity originating from other brain structures. In particular, PMC represents the main source of the excitatory input for striatum and STN. Moreover, its activity encodes competing motor responses modulated by BG; indeed, in the initial stages of training, when the BG is not able to facilitate a response, random noise in PMC is required to allow the network to select a random response.

### 4.3. Neurons and synapses

In order to make everything as simple as possible, but not simpler, we used one type of neuron model in our BG network. The goal is to realize a baseline implementation, and better characterize the neural network in a future work with respect to experimental and physiological data. All neurons in the network were realized using the leaky-integrate-fire neuron (LIF) model with exponential function-shaped postsynaptic conductance [30], which has been shown to be consistent with experimental data on the parameters characterizing in vivo-like activity of cortical neurons [31] and recently being adopted in several models of the BG using spiking neurons [32, 33, 34, 35, 36]. While LIF neuron model may appear simple, we can vary its inputs and parameters to fit different input-output firing rate relationships (as we have done). The size of the layers have been chosen in accordance with Frank’s models [3, 9], tending to simulation efficacy at the expense of biological details and the ability to capture neuronal dynamics (synchronization or oscillations). Also, the network parameters are obtained from the associated biological values through a process of normalization. The neural parameters are given in the Table S2.

The projections from input to striatum and to the PMC are plastic. All remaining connections of the direct, indirect and hyperdirect pathway and the recurrent interactions within BG are static conductance-based synapses determined to assure the fulfillment of a set of functional restrictions by a robustness analysis and are in line with those used in Frank’s models [3, 9]. All synaptic connectivity parameters are specified in Table S3.

### 4.4. Learning rule

The projections from the cortical input layer to the BG (striatum) and to the cortex (PMC) are learnable by two different learning mechanisms. However, the learning rule is identical for each connection, independent of the pathway or DA receptor or column.

For cortico-striatal learning, we have used a reinforcement learning version of Leabra, the rule proposed by Frank [9], which is more biologically plausible than standard error backpropagation. To simulate feedback effects, the learning algorithm involve two distinct phases: “minus” and “plus” phase. In the minus phase, the network settles into activity states based on input stimuli and its synaptic weights, ultimately choosing a response. In the plus phase, the network resettles in the same manner, with the only difference being a change in simulated DA release (i.e: SNc activity) depending on the selected response: DAergic neurons increase from tonic to phasic (i.e: high) level of activity, which means that the unit firing rate goes from 0.5 to 1.0 for a correct response, whereas an incorrect response causes a decrease from tonic to zero level of activity, meaning from rate of 0.5 to 0.

Specifically, the rule uses a combination of error-driven and Hebbian learning. The error-driven component computes a simple difference of a pre- and post-synaptic activation product across the two phases. For Hebbian learning, we use the same learning rule used in competitive learning. The error-driven and Hebbian learning components are combined additively at each connection to produce a net weight change.

The equation for the Hebbian weight changes adopted is:

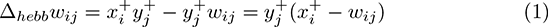

where *x* is the presynaptic activation, *y* the postsynaptic activation and −/ + represents minus/plus phase, respectively. The equation for error-driven component is:

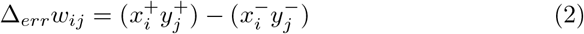

which is subject to a soft-weight bounding to keep within the 0–1 range:

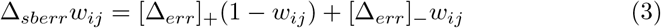

The two terms are then combined additively:

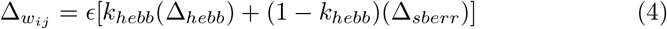

where *ϵ* represents the learning rate parameter and *k*_*hebb*_ is a parameter that controls the associated proportion of the two types of learning.

In our case, the plasticity depends only on the reward value. Therefore the mechanism is a reward value based learning that triggers plasticity regardless of what was expected. Note that an explicit supervised training signal is never presented; striatal connection weights change depending on the difference between activity states in the minus and plus phases, which only differ due to phasic changes in DA. The striatum learns over time which responses to facilitate and which to suppress in the context of incoming sensory input, but whether there is LTP or LTD, the model is completely in the dark. LTP (or LTD) occurs if neuronal activity is increased (decreased) with respect to the minus phase. In addition, the PMC itself learns to favor a given response for a particular input stimulus, via Hebbian learning from the input layer (Equation 1). The parameters regarding the learning mechanism are in Table S4.

Thus, the BG initially learns which response to gate via phasic changes in DA ensuing from random cortical responses, and then this learning transfers to the cortex once it starts to select the correct response more often than not. This implements the idea that the BG modulates the gating of responses that are selected in the cortex. Both learning rules do not include an explicit maximum weight value (we omitted because it is irrelevant for the type of analysis of the present work), but include a normalization of each term considered.

### 4.5. Data Analysis

All data were processed using custom scripts in Python and standard statistical analyses. The significance level for statistical tests was *α* = 0.05. The F-test adopted is one-way ANOVA. Unless stated otherwise, the simulations are repeated for 30 networks for each experimental group. We considered performance of 30 network to be robust to different instantiations of network parameters initialization. As the computational model does not include the cerebellum, parameters such as timing, velocity, latency or jitter have no significance to be measured. On the other side, we provide some qualitative parameters which could be useful to compare results with experimental or behavioral studies involving gradual DA depletion.

## 5. Task description

In all the experiments we simulated a task to test the capacity of the network to learn a match between stimuli and actions, to learn a novel motor program. Motor learning allows to learn and refine new skills through practice, as they are associated with long-lasting neuronal changes. Motor learning processes strictly depend on the structural integrity and functional activity of the cortico-striatal loop (discussed in Section 2) and cerebellum. Specifically, skilled movement involves combining smaller elements of the movement in a particular order with certain timing; in other words, a motor skill is the ability to perform complex sequences of movements quickly and accurately. While other brain structures are thought to be involved in motor planning and coordination, we consider BG to be crucial in learning the cortico-striatal association to accurately control the release of an action in the cortex and the suppression of all the competing ones (discussed in Section 2). So, the BG is part of motor planning, but do not implement this function. For this reason, we adopt a sequenced simulated motor task, but our BG network do not memorize sequence production information.

The network has four possible stimuli and four possible actions. The stimulus presented in each trial is presented for the whole duration of the minus phase, that is dynamic, and follows the sequence shown in Figure 3, but we assured that the network shows the same learning behavior also if the stimulus was selected randomly. Full success is achieved if 40 correct answers are given following each other, which mean 10 repetitions of the task (namely, epochs).

**Figure 3:**
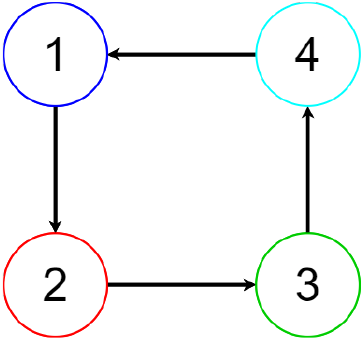
Simulated motor task. Given an input signal, the network has to learn and select the most appropriate motor command to fulfill the task, in line with the hypothesis of complex motor behavior composed of elementary motor responses. The simulated task involve four different motor responses associated to different stimuli. The network is considered to have properly performed the task when the four responses are given for the matching input to generate the motor program.

Interestingly, this level of abstraction of the motor task allows us to include among the possible tasks considered in the simulations many different experiments currently carried out on different species and to easily address results to experimental scope. Indeed, sequence learning paradigm has been used extensively to study motor skill acquisition and, on the other hand, as the task we modeled is high level, any behavioral complex motor task may be reduced to a set of simplest motor elements. Motor learning tasks of this type are commonly used in experimental studies to analyse motor skill acquisition and long-term exercise association with improved performance. For example, in mice [23, 24]; in humans, hand-writing tasks [37, 38], physical exercise [39], finger button presses [40, 41], joystick movements [42], and others. Motor learning performance (improvement or degradation) following the repetition of specific commands is then evaluated behaviorally in terms of movement speed, accuracy, latency, total time, distance traveled, or similar.

## 6. Results

Frank’s model [3] predicts that DA is critical for learning and executing responses, particularly phasic changes and a large dynamic range; therefore, a reduced dynamic range of DA explains Parkinson. Specifically, decreased levels of DA would lead to spared NoGo learning and impaired Go learning, as it depends on DA bursts. In our experiments, we stand on the shoulder of giants and present a novel computational exploration of the role of DA deficiency in motor learning within the overall BG circuitry. We used numerical simulations of the BG network with spiking neurons to understand how different levels of DA depletion shape motor learning, especially the prodrome. In the model, we systematically varied the DA level and studied how gradual reductions affect the behavior with respect to healthy condition. Here, we set the DA level from 1.0 to 0.5 to tune the model into healthy and PD-symptomatic conditions respectively, more details further in the text.

### 6.1. Simulations 1a-c: healthy conditions

The job of the network (see Figure 2) is to select either R1 or R2 or R3 or R4, depending on the sensory input. At the beginning of each trial, incoming stimuli directly activate a response in the MC. However, these direct connections are not strong enough to elicit a robust response in and of themselves; they also require bottom-up support from the thalamus. The job of the BG is to integrate stimulus input with the dominant response selected by the MC, and on the basis of what it has learned in past experience, either facilitate (Go) or suppress (No-Go) that response. Within the overall thalamocortical circuit, there are two parallel subloops that separately modulates the four responses, allowing the BG to selectively gate one response, while continuing to suppress the others. In this regard, the striatum is functionally divided into two subpopulations. The four columns on the left are “Go” units for the four potential responses and simulate D1R. The four columns on the right are “NoGo” units and simulate D2R. Thus, the eight columns in the striatum represent “Go–R1”, “Go–R2”, “Go–R3”, “Go–R4”, “NoGo–R1”, “NoGo–R2”, “NoGo–R3”, “NoGo–R4”. The Go columns project only to the corresponding column in the GPi (direct pathway), and the NoGo columns to the GPe (indirect pathway). The GPe columns inhibit the associated column in GPi, so that striatal Go and NoGo activity have opposing effects on the GPi. Finally, each column in the GPi tonically inhibits the associated column of the thalamus, which is reciprocally connected to the MC (PMC and M1). Thus, if Go activity is stronger than NoGo activity for R1, the left column of the GPi will be inhibited, removing tonic inhibition (i.e., disinhibiting) of the corresponding thalamic unit, and facilitating its execution in the MC.

At the beginning of training, random activity within the network causes the selection of a random answer. During learning the activity within each striatal column enables different Go and NoGo representations to develop for various stimulus configurations. In particular, the DA mechanism modulates striatal learning, since increases in DA during positive feedback lead to reinforcing the selected response, whereas decreases in DA during negative feedback lead to learning not to select that response. When the network has strengthened the correct Go/NoGo connections and is able to steadily complete the task, learning is completed.

In this regard, the simulations have been carried out for 300 epochs (i.e: 1200 trials) in order to test the steady-state behavior. By considering the whole simulation time (refer to Figure 4), we consider early stage of learning from epoch 0 to 30, learning stage from 30 to 80 and completed or after learning from 80 on. The early stage is assessed as the average error falls below chance (50%); the completion of learning has been evaluated as when the average error remains below 5% for 10 consecutive epochs, defining the learning time.

**Figure 4:**
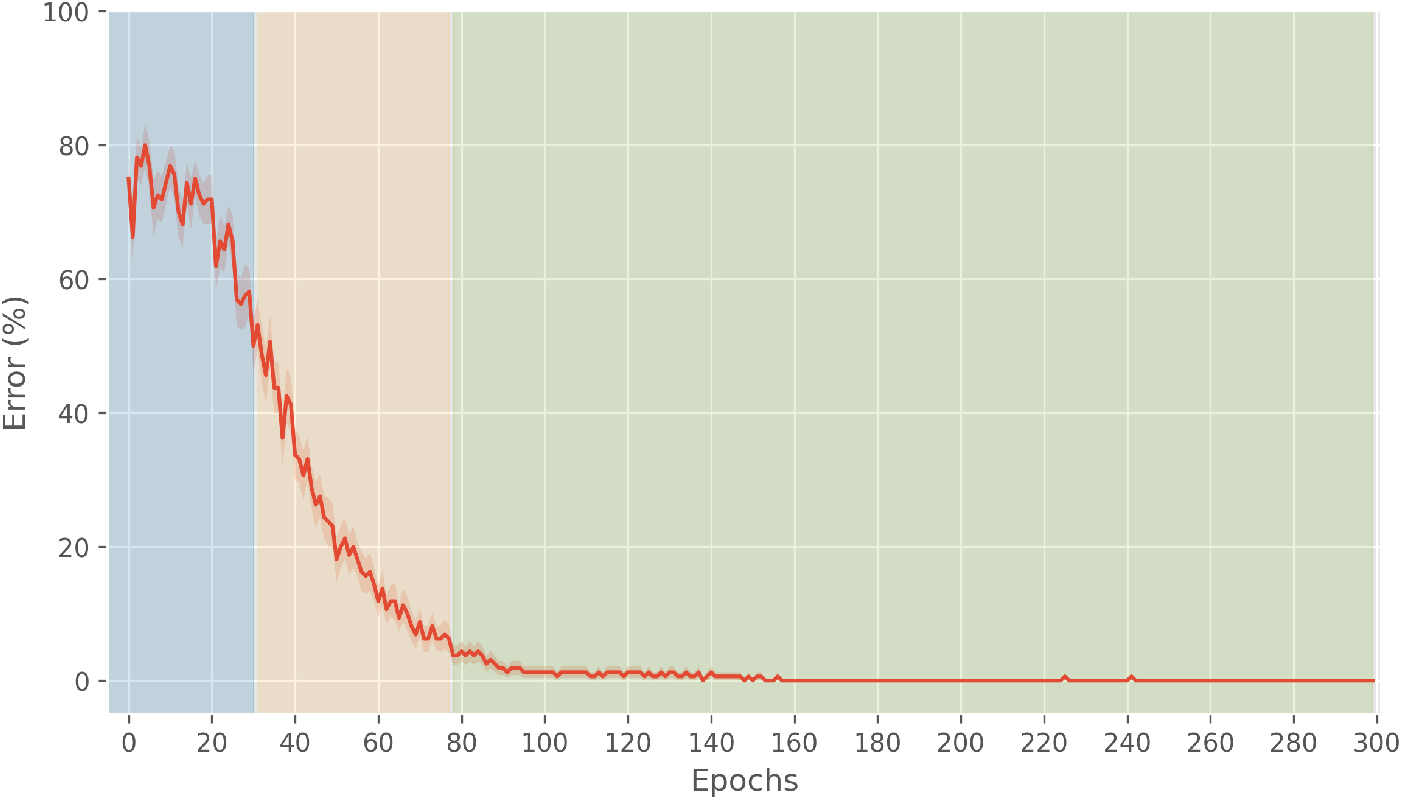
Learning error curve. In blue, the early stage of learning, namely early phase; in yellow, learning phase; in green, after learning phase

#### 6.1.1. Simulation 1a: early phase

In the first epochs, the GPi and GPe are massively activated due to the background neural activity; STN activity is influenced by the background neural activity and cortical afferents, exhibiting a decay of activation within the trial simulation; PMC is strongly dependent on the random background activity and subsequently M1 activation is random. At the beginning, striatal weights, stimulated by cortical input, are not strong enough to disinhibit a column in the GPi to select a response, through the thalamus. Indeed, the thalamus is not active up to around the end of the early phase. Actually, activation in the striatum is scarce, since they are triggered by the combined effects of tonic DA, cortical input and PMC.

The analysis of raster plots and weights (here omitted) showed that this behavior starts to change towards the end of the early phase and hints of learning are: neurons in the Go-striatum leading to the selection of the correct response being enhanced, those in the Go-striatum leading to the selection of a wrong response being weakened, those in the NoGo-striatum leading to the selection of a wrong response being enhanced and those in the NoGo-striatum leading to the selection of the correct response being weakened. This activation in the striatum causes changing behavior in its afferents: some neurons are inhibited in the GPi corresponding to a correct response, as an effect of behavioral change in striatum and STN (through GPe). In this phase, Go pathway activation, albeit increasing, is not enough to select the correct response. It is remarkable that striatal neurons, although receiving the same stimuli, exhibit different activity: due to the inclusion of striatal interneurons, some neurons are not firing and some exhibit different firing rates from others.

#### 6.1.2. Simulation 1b: learning phase

In this phase, striatal weights are consistent to inhibit the GPi column corresponding to at least one response, and lead to the disinhibition of the corresponding neuron in the thalamus. The activation of the thalamus results in a visibly appreciable enhancement of cortical activation in PMC, now provoked by Go pathway, through the thalamus, and not by random cortical activity.

Striatal activation is enhanced with respect to the previous phase, meaning that learning is proceeding and the connection with the cortical input is being strengthened, although the network indecision is still reflected in activation for the other responses, because DA release and interneurons connection are fixed. At the end of the learning phase, as it is defined, striatal weights have become capable of systematically facilitating the four responses.

#### 6.1.3. Simulation 1c: after learning phase

Learning can be considered completed for the motor task, as the network has learned the correct SR associations for all the responses and the behavior is constant in time. For each stimulus presented, the inhibition of the associated column of the GPi is substantial and robust, as well as the activity in the thalamus and PMC, because Go pathway activation is massive for the correct response and inhibited for the other responses.

### 6.2. Simulations 2a-b: pathological conditions

PD is a BG dysfunction associated with a decrease in the production of DA in the SNc, which affects the normal behavior of the BG. Aggregation of misfolded proteins has been suggested to be the characteristic feature of the neurodegenerative process, despite it is not clear whether Lewy bodies (LBs) are causes or scars of neurodegeneration. What is clear is that LBs formation in DAergic neurons slow down DA metabolism, slowly toning down the amount of DA in the striatum. From this point, the aim is that of simulating the reduction of the amount of DA in the striatum.

We consider that the reduction of DA can be achieved implementing different kind of lesions in a computational model. In particular, (i) to cut DAergic neuron connection, (ii) decrease weight synapse connection between SNc and striatum, and (iii) decrease DAergic neurons firing rate. Simulating PD by lesioning the network in any of the three cases means to reduce dynamic range of the DA signal. Dynamic range is critical for learning appropriate Go/NoGo representations from error feedback, as network weights are adjusted on the basis of difference in activity states in the two phases of network settling. Because tonic DA levels are low, PD networks have an overall propensity for NoGo learning, but the dynamic range of phasic dips during negative feedback is reduced. Go learning is degraded because limited amounts of available DA reduce the potency of phasic bursts, activating less of a Go representation during positive feedback.

The first type of lesion, adopted by Frank [3], is the less similar to the neurophysiology and the sharpest in terms of damage given the size of the SNc in the model. The second type impacts the firing rate of DAergic neurons indirectly and reduces the amount of DA which reaches the striatum, whereas third one acts directly on the firing rate. Also, if a neuron stops firing or it does not affect postsynaptic terminals anymore, it can be considered dead.

We made a first investigation exploring all the three lesions listed above, in order to evaluate the impact of the degree of a damaged parameter over the network behavior. Both the alteration of striatal weights and firing rate are interesting and employed in other computational studies (i.e: [33]), but we considered manipulating synapses with purpose to stand at a microscopic scale and out of context with this work. Moreover, the rate modulation allows a fine-granularity manipulation suitable for studying different stages of the disease. For these reasons, we simulated SNc lesion in our network by setting to different values the rate of the Poisson process driving the SNc unit.

The experimentation has been set up taking a look at clinical findings. Currently, the literature still falls far short of a uniform and well-defined view of PD disease development, neither of diagnosis and treatment.

Ugrumov [43] claims that according to the last classification, three stages are distinguished in PD pathogenesis: (i) the preclinical stage — from the onset of neurodegeneration to the appearance of nonmotor symptoms, (ii) the prodromal stage — from the onset of nonmotor symptoms to the appearance of motor symptoms, bradykinesia, rigidity, rest tremor, (iii) clinical stage — from the appearance of motor symptoms to death. PD develops up to 30 years at the preclinical stage without manifestations of motor disorders. They appear after a “threshold” degradation of the nigrostriatal DAergic system at a loss of 70% DA in the striatum and 50-60% DAergic neurons in the SN (Figure 5).

**Figure 5:**
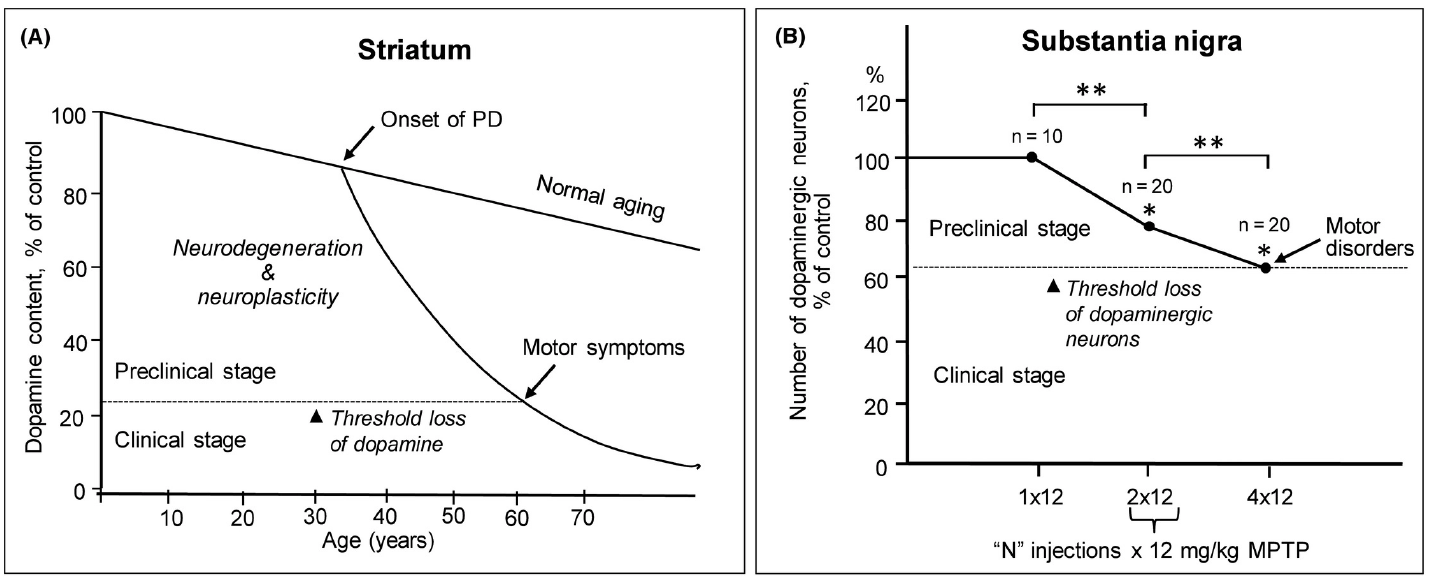
Schematic representation of the pathogenesis of PD in patients and its reproducing in animal models. A, The manifestation of progressive degradation of the nigrostriatal DAergic system in patients with PD as a loss of DA in the striatum to a threshold level, which is associated with the onset of motor disorders; B, Progressive loss of DAergic neurons in the substantia nigra when modeling preclinical and clinical stages of PD in mice by subcutaneous injections. Reproduced from [43, Figure 1A,D].

Grosch et al. [44] claim that the loss of DAergic striatal nerve terminals at motor symptoms onset is rather difficult to determine, reporting a comparison between different studies performing regression analysis to the onset of motor symptoms to estimate the proportion of lost striatal and putaminal DAergic terminals, which goes from 39% to 51% for striatum and 51% to 71% for the putamen.

Cheng et al. [45] suggest that at the time of first diagnosis of PD, only 30% or so of DAergic neurons have been lost and 50-60% of their axon terminals. Moreover, they conclude that at the time of motor symptom onset the extent of loss of striatal DAergic markers exceeds that of SN DA neurons, in accordance with Ugrumov. This conclusion is consistent with observations that, at the time of death, depending on disease duration, while there has been 60-80% loss of SN DA neurons there has been a much more profound loss of striatal or putaminal DAergic markers.

Lastly, findings indicate that degeneration of DAergic neurons in PD is not total even after many years of illness [46] and at the time of death.

Then, which stages should be investigated? The low efficacy of treating patients is due to the late diagnosis and start of therapy, after the degeneration of most specific neurons and depletion of neuroplasticity. It is believed that the development of early diagnosis (ED) and preventive treatment will delay the onset of specific and severe symptoms.

The current methodology for the development of ED is based on searching biomarkers, such as premotor symptoms. In this direction, handwriting is currently considered a potential powerful marker for the ED of PD, with the purpose of building tailored intelligent systems to support clinicians based on machine learning techniques [47, 48, 38, 49]. Handwriting, indeed, is one of the abilities that is affected by PD; because of that, researchers investigate the possibility of using handwriting alterations, caused by the disease, as diagnostic signs, expression of small entity of SN lesion, in the early stage of the disease, before the onset of evident symptoms. This hypothesis is well-founded and based on the fact that in the clinical practice there is already employed a writing exam for the diagnosis of neuropsychological deficits, employed also for PD, which comprises asking the patient to create a series of specific drawings. Despite this, though, no specific writing test has been approved or indicated in the guidelines of clinical practice for PD diagnosis. Nevertheless, these studies show that, even when the clinical diagnosis is at an early stage (based on UPDRS) and subjects do not show obvious motor deficits, the analysis of the dynamic characteristics (such as speed, acceleration, fluence) associated with a fine motor task, such as the generation of writing, already allows to highlight some differences between healthy subjects and patients. PD patients are also known to have motor learning difficulties [20]. Consequently, the execution of a new motor task is more deficient than the execution of a known motor task. Furthermore, it has been shown that there are similarities between the dynamic characteristics of the writing movements of healthy subjects when they produce a motor task new to them and those measured in subjects who have a PD in the early stage [49].

Our experimentation goes along with this research area, as it may be of high interest to investigate the learning deficits shown by the model for different stages of DAergic degeneration, and to gain some indications on the possibility of developing a protocol that would be simple to perform, inexpensive and non-invasive and, thus, allow for early detection of the disease.

#### 6.2.1. Simulation 2a: PD pathogenesis

The damaged networks have been simulated for 300 epochs (i.e: 1200 trials), to be comparable with the healthy networks. The lesion range is 0.8, 0.7, 0.6, 0.5, 0.3, where 1.0 is healthy and the lesion is inflicted at the beginning of training.

Given the results obtained for the healthy networks (see Figure 4) and the clinic remarks, we are interested in analysing the behavior of preclinical and prodromal PD in the model. Claiming the incapacity to establish a complete matching with the physiology, we may consider the range 0.8, 0.7, 0.6 as a simulation of preclinical stage and from 0.5 of motor symptoms and clinical stage.

Error curves are shown in Figure 6. It is clearly visible a change of shape in 6e, 6f, as 0.5 and 0.3 are a degree of damage which produce evident misbehavior. Consistent with the prediction of Frank’s model [3, 9], the model suggests that persistent heavy DA depletion affects the steady-state of the BG networks, which were impaired, and also results in stronger activity along the indirect pathway.

**Figure 6:**
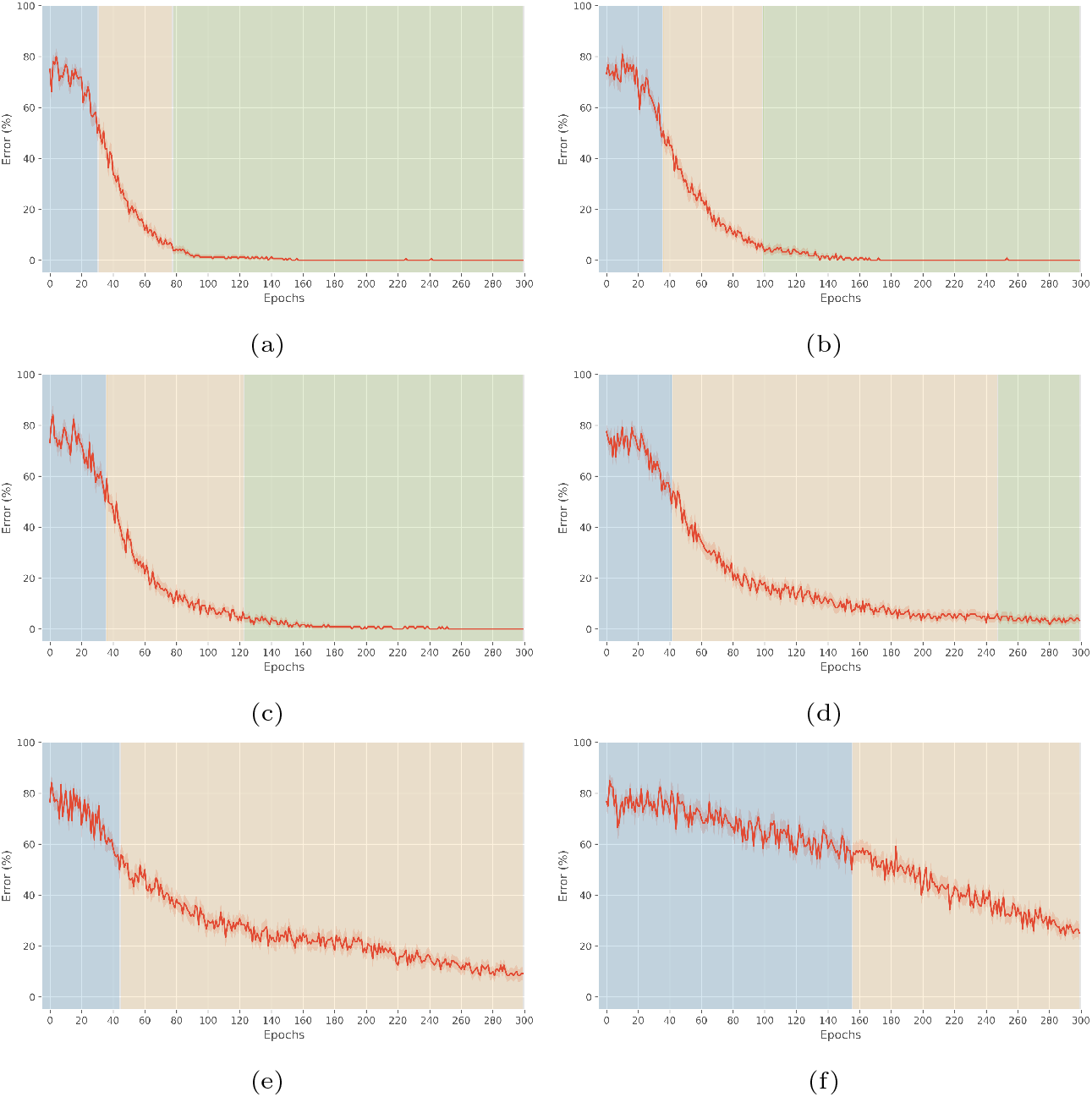
The correspondence between DA depletion level and the value of firing rate: (a) 1.0: 1000.0 spikes/s (b) 0.8: 800.0 spikes/s, (c) 0.7: 700.0 spikes/s, (d) 0.6: 600.0 spikes/s, (e) 0.5: 500.0 spikes/s, (f) 0.3: 300.0 spikes/s. In blue, early phase, in yellow, learning phase, in green, after learning phase, calculated for each condition respectively.

Also, the learning phase (yellow area in Figure 6), as we have defined it, shows an increasing duration with the increase of the entity of the lesion.

Focusing on the learning phase of the healthy condition, the learning curve is statistically different between healthy and damaged networks, also in case of small lesion (Figure 7). The qualitative parameters calculated are: percent error in the first 5 epochs considering early phase of learning, i.e: 0-5 (Figure 8a), percent error in the first 5 epochs considering learning phase of healthy condition, i.e: 30-35 (Figure 8b), percent error in the last 5 epochs considering learning phase of healthy condition, i.e: 75-80 (Figure 8c), learning time (Figure 8d), defined as the epoch in which the error remains below 5% for 10 consecutive epochs, slope (Figure 8e) and intercept (Figure 8f) of the motor task learning curves in Figure 7.

**Figure 7:**
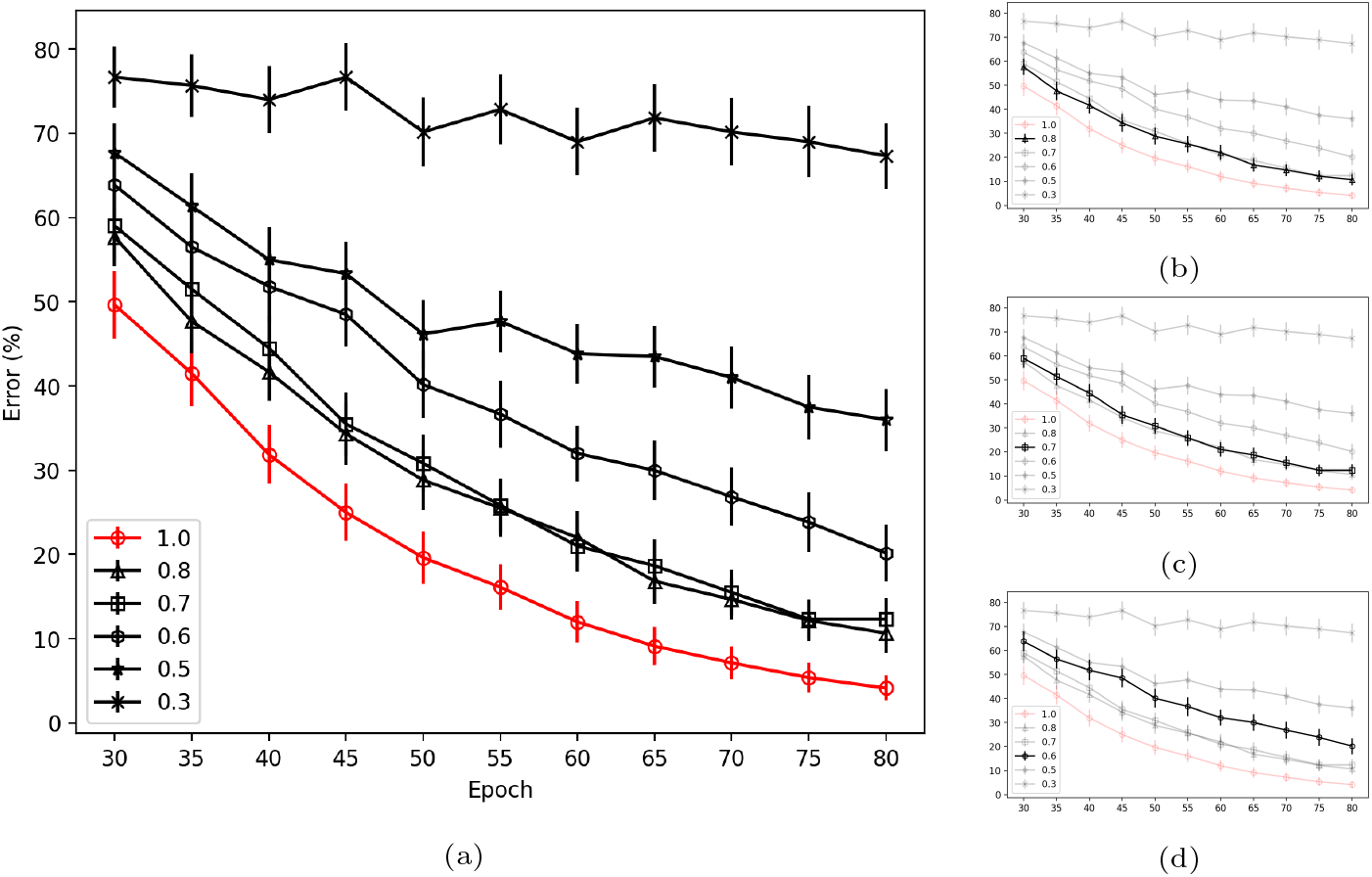
Motor task learning curves, averaged over 30 networks for each condition. Intact: Full BG model with direct and indirect pathways modulated by phasic changes in simulated DA during error feedback, corresponds to 1.0; PD: simulated Parkinson’s disease, modeled by lesioning firing rate of DAergic units in SNc, range 0.8, 0.7, 0.6, 0.5, 0.3. PD networks are impaired at learning the task, due to impoverished phasic changes in DA in response to feedback. Impairment occurs also for less damaged networks (0.8, 0.7). In contrast, intact models can use response-specific Go and No-Go representations that develop over training in order to more selectively facilitate and suppress responses. Learning curve emphasized for simulated preclinical PD in 7b, 7c and 7d.

**Figure 8:**
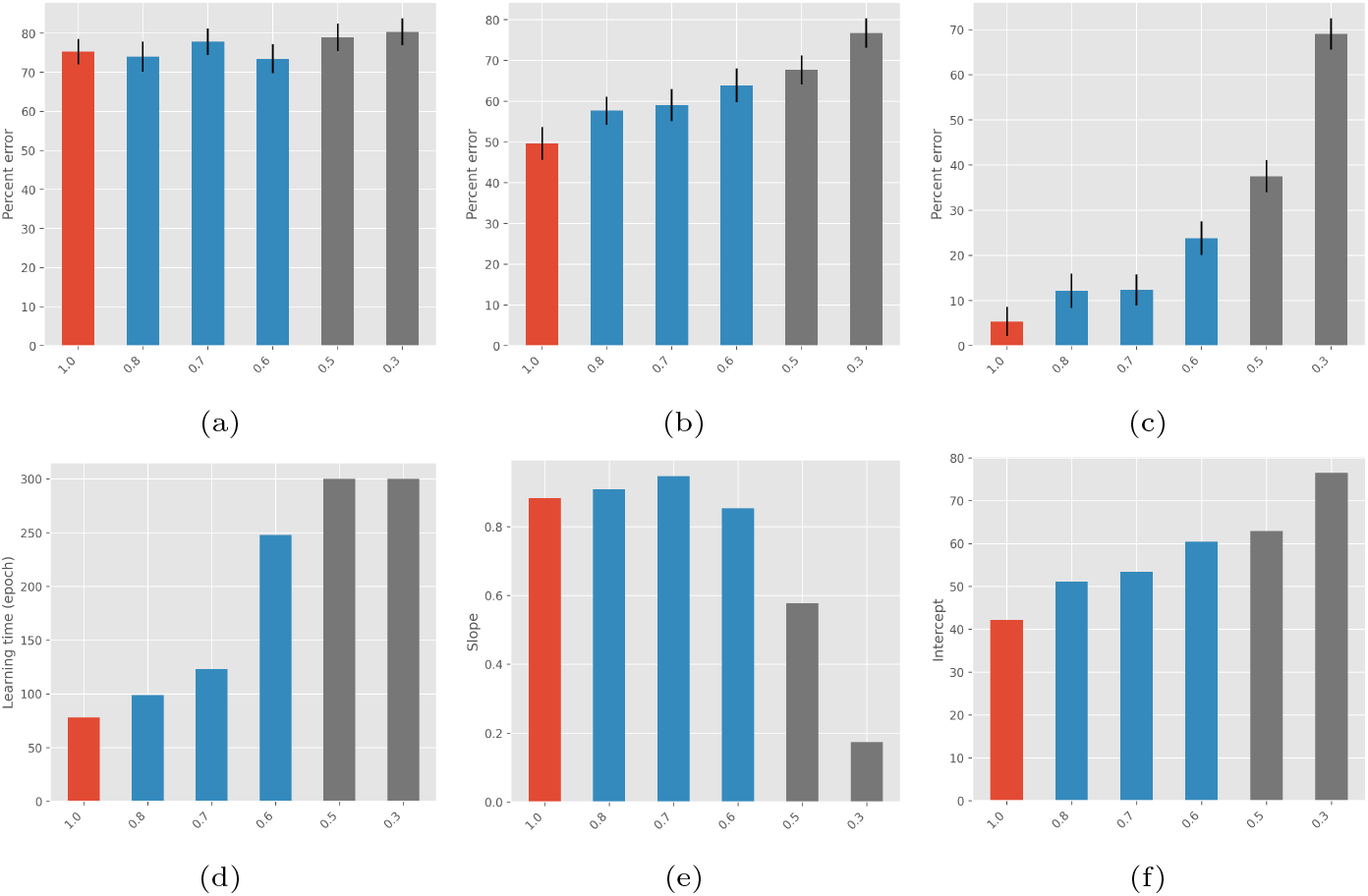
Qualitative parameters, ranging from healthy (1.0, red) through preclinical PD (0.8, 0.7, 0.6, blue) to clinical PD (0.5, 0.3, grey) stages: (a) First 5 epochs average error of early phase (b) First 5 epochs average error of learning phase (c) Last 5 epochs average error of learning phase (d) Learning times (e) Slope (absolute values) (f) Intercept.

Intact networks successfully learned the motor task, scoring 95% optimal responding after 78 epochs (312 trials) of training (Figure 4). “Parkinson” networks were impaired, only scoring 87%, 87%, 76%, 62%, and 31% optimal responding respectively (Figures 7, 8c). Statistical analysis indicated that this difference was highly significant [F(5,174) = 101.78, p *<* .001]; specifically, there was a statistically significant difference in score between simulated preclinical PD and healthy networks [F(3,116) = 12.87, p *<* .001]. Also, comparing the performance of networks with the smallest degree of lesion with healthy data, they are significantly different [F(1,58) = 7.46, p = .008]. In this regard, learning time trend increases exponentially with the degree of DA loss (Figure 8d). In contrast, there is no statistical difference in the performance over first epochs of training [F(5,174) = 0.61, p = .69] (Figure 8a), coherently with the model design, whereas different performance in the first epochs of learning phase is highly significant [F(5,174) = 5.81, p *<* .001] (Figure 8b).

The regression analysis of the learning curves shows that while the steepness (Figure 8e) does not change significantly between healthy and preclinical PD (instead it decreases, as its magnitude decreases, for clinical PD); in contrast intercept (Figure 8f) consistently reflects the trend of the performance in early learning phase (Figure 8b). This result coherently predicts that preclinical PD networks were able to learn the motor task similarly to healthy networks but they required more time to reduce the error and carry on their learning process, given that their learning phase has a different (growing) starting error. In the case of clinical PD, learning does not reach a steady-state within the time of simulation; this is predicted by the highly reduced slope of the curves.

Lastly, results show very close curves and parameters for 0.8 and 0.7 lesioned networks, which are the most interesting as they represent the smallest lesion inflicted to the model, but both are statistically different from healthy networks.

#### 6.2.2. Simulation 2b: further simulations

For consistency, simulations involving the insertion of the lesion at half (i.e: during learning phase) and after the completion of learning (i.e: after learning phase) were done, for the range of 0.7, 0.5, 0.3 (not shown). By damaging the networks at half learning, the behavior is slightly deteriorated with respect to healthy networks, similarly whether the entity of lesion is smaller or larger, but with an increasing SEM. By damaging the networks after learning, as BG response is consolidated by the strengthening of cortico-striatal synapses, we report that there is no statistically significant difference with the healthy condition. It is consistent with the findings that PD patients are able to continue performing already learned motor tasks, although not smoothly. In fact, parkinsonian tremor tends to decrease or disappear with the performance of voluntary movements [50].

## 7. Discussion

This work presents a theoretical basis for motor learning functions and alterations of the BG. A neural network mechanistic model incorporating known biological constraints provides an insight into the pathogenesis of PD and motor learning deficits.

Consistent with Frank’s model, a key aspect of the model is that phasic changes in DA during error feedback are critical for the implicit learning of SR associations. In addition, the model incorporates the inhibitory effect of interneurons in the striatum and the global NoGo of STN, as well as direct, indirect and hyperdirect pathway and DAergic system. Despite the expansion of the model, simulations in healthy conditions confirm model predictions provided by Frank.

The model has been used to provide an innovative account of motor learning deficits in PD conditions. Simulated Parkinsonism has been obtained by reducing the amount of DA in the model, through the manipulation of firing rate in SNc and, thus, of its modulatory effects on Go and No-Go representations, produced qualitatively similar results to those obtained by Frank and supports an investigation of the preclinical and prodromal stage of the illness.

Damaging the network in varying degrees has resulted in the validation of the hypothesis that the BG play a key role during learning. Furthermore, the deteriorated performance in handwriting tasks, found in the experimental studies on PD patients, is echoed in the deteriorated learning behavior of the networks when a DA lesion is inflicted. Our findings are that some symptoms, which may be addressed as motor learning symptoms, actually arise in the simulated Parkinsonian networks before the evident motor symptoms, with minor damage to the DAergic system. Specifically, the learning curves show that a 20% DA loss already causes a significant decline of the performance with respect to the healthy networks in learning a novel motor skill, thereby confirming that learning symptoms arise at early stage of the disease. In other words, we can claim that, with 20% DA loss in the networks, learning is slowed down and the dynamic properties are different with respect to the healthy case. Novel predictions from this model have to be subsequently confirmed in Parkinson patients.

This result is in line with handwriting analysis and handwritten pattern recognition findings (reviewed above) and takes a step further. The other techniques have proved that discriminating between unhealthy and healthy subjects on the basis of simple and easy-to-perform handwriting tasks is possible, but they are based on data from patients already diagnosed with PD. Instead, this work proves that there may exist abnormalities of motor learning process, due to alterations in the dynamics of BG functionality, which do not yet involve the presence of symptoms that normally lead to clinical diagnosis.

Furthermore, considering that the feature of parkinsonian movements is that PD patients are not able to properly develop a motor plan, made of properly synchronized smaller elementary movements to execute a more complex sequence of movements in such a way as to minimize energy consumption and to execute a motor task smoothly [38], the performed movement will be composed of many stop-an-go, interspersed with properly synchronized movements whose extent decreases as the size of the lesion increases. In other words, PD patients are not able to learn the complex movement and they segment it into pieces. Taking as an example handwriting, in the early stage of the disease, the feature of the deteriorated motor plan does not concern the shape of the handwriting, rather the dynamics and the variations in acceleration. Specifically, the work proves that these variations in acceleration concern the dynamics of learning a novel motor task and are not found in the execution of already learned tasks.

Thereby, as motor learning results in smooth handwriting, in PD patients this process is not complete, and the handwriting movements are jerky; this feature can be studied by analysing velocity and acceleration profiles. What emerges from this study is that a higher value of normalized jerk, the time-derivative of acceleration, and abrupt changes in movement velocity and acceleration during the dynamics of novel motor learning, may result also for patients not yet diagnosed PD with respect to healthy subjects, while so far it has been proved only for patients already diagnosed with PD. Then, the work reveals that while many features in the literature concern the shape of the handwriting, actually it is not as important as the dynamic producing that shape, which may be measured by velocity and acceleration, for instance.

In summary, results obtained provide a working computational feedback for the hypothesis of deteriorated behavior in motor learning task, as handwriting, for preclinical PD, that can be further tested experimentally and behaviorally at different scales and more granularity. Also, findings give an indication that subjects should be tested on the execution of novel and relatively complex motor tasks and that parameters of the dynamics producing the movement in the learning phase should be evaluated to differentiate between healthy and non healthy undiagnosed subjects for an ED, but also for assessing the stage of the disease or for the definition of novel rehabilitation protocols and other treatments.

## 8. Conclusions

The computational study of neurodegeneration in the BG revealed that motor skill learning is affected by DA depletion and even small loss of DA, which may be associated to early stage of PD, interestingly alters the process. This result could shed some light on the discovery of biomarkers of the disease in preclinical and preclinical PD, currently unidentified. This work should provide direction for further experimental studies investigating both learning curves and biological and behavioral parameters in learning new motor skills and the relationship between the learning profiles and the other parameters. The characterization of such correlation would represent a significant stepping stone towards the research of the issue.

## Acknowledgments

This research was supported by the European Union’s Horizon 2020 Framework Programme for Research and Innovation under H2020-EIC-FETPROACT-2019 Grant Agreement 951910 (MAIA) to I.S.

## Conflicts of Interest

The authors declare no conflict of interest.

## Supplementary Material

**Table S1:**
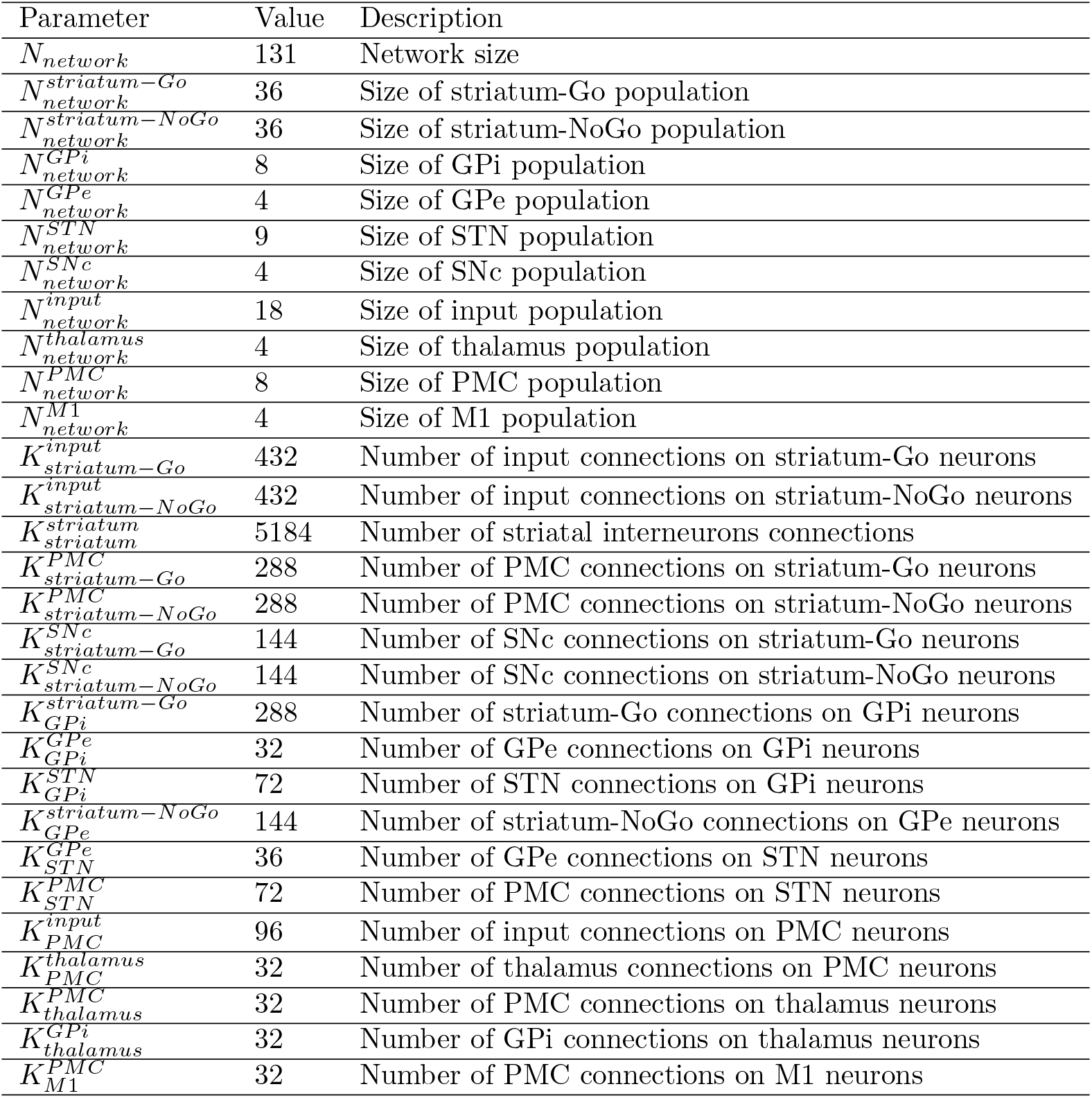
Network and connection parameters

**Table S2:** Neuron parameters

| Parameter | Value | Description |
| --- | --- | --- |
| $C_m$ | 2.0 | Membrane capacitance |
| $g_L$ | 0.2 | Leaky conductance |
| $E_L$ | -70.0 | Leak reversal potential |
| $V_{reset}$ | -70.0 | Reset potential of the membrane |
| $V_{th}$ | -40.0 | Spike threshold |
| $\tau_{syn\_ex}$ | 0.5 | Rise time of excitatory synaptic conductance |
| $\tau_{syn\_in}$ | 10.0 | Rise time of inhibitory synaptic conductance |
| $E_{ex}$ | 0.0 | Excitatory reversal potential |
| $E_{in}$ | -85.0 | Inhibitory reversal potential |
| $I_e$ | 0.0 | Constant input current |
| $t_{ref}$ | 1.0 | Duration of refractory period |

**Table S3:**
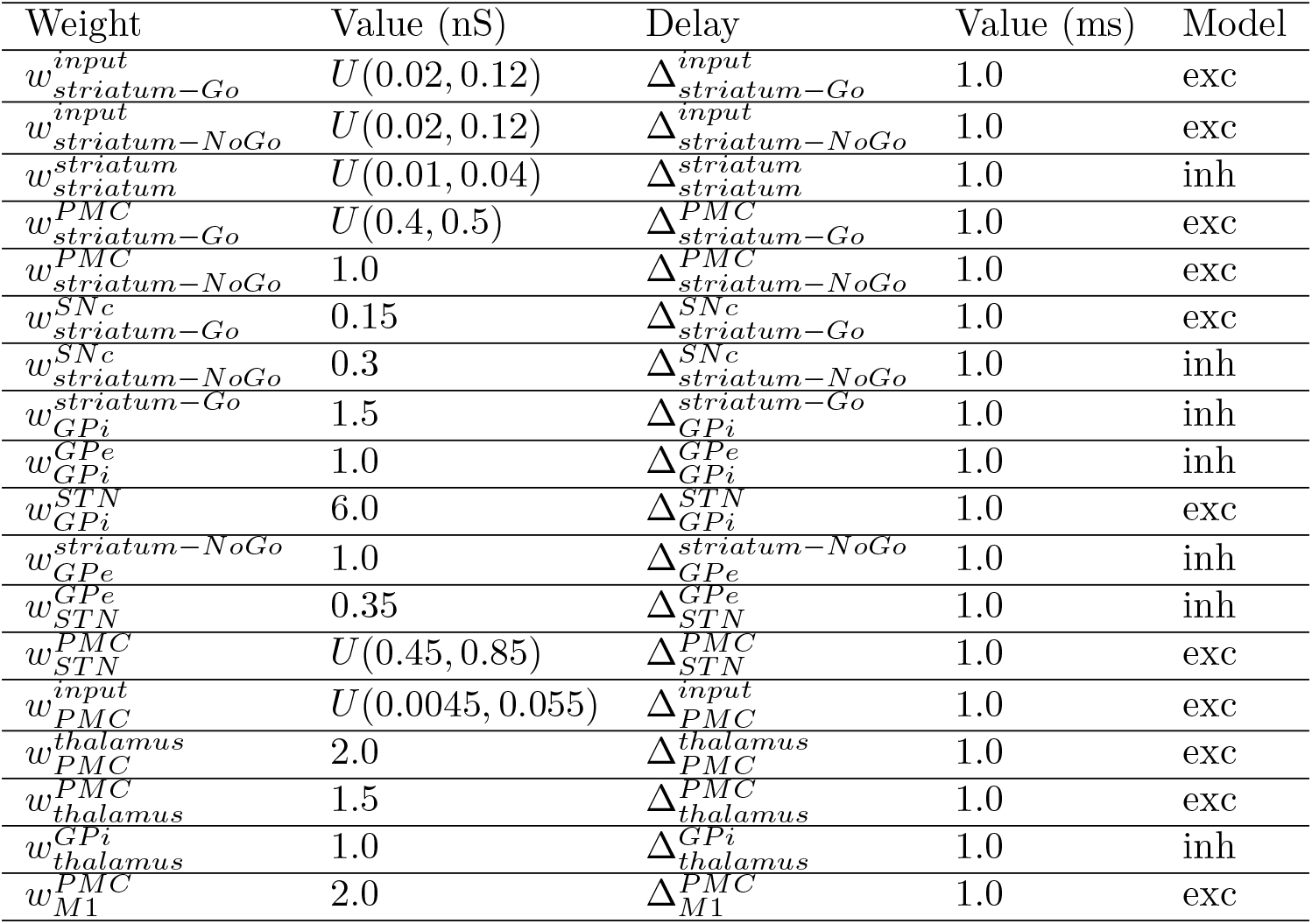
Synaptic parameters in healthy condition

**Table S4:** Learning rule parameters for the connection between the input cortex and the BG (striatum) and associated simulation parameters

| Parameter | Value | Description |
| --- | --- | --- |
| $Tw^-$ | Variable | Duration of the minus phase |
| $step^-$ | $2ms$ | Step size of simulation |
| $Tw_{min}^-$ | $400ms$ | Minimum time of simulation |
| $Tw_{max}^-$ | $800ms$ | Maximum time of simulation |
| $Tw_{sp}^-$ | $60ms$ | Time window for evaluating firing activity |
| $Tw_{th}^-$ | 0.25 | Threshold firing rate as $n_{sp} \div Tw_{sp}^-$ |
| $Tw^+$ | $200ms$ | Duration of the plus phase |
| $Tw_{cor-str}$ | $60ms$ | Time window for striatal learning |
| $Tw_{cor-cor}$ | $60ms$ | Time window for cortical learning |
| $\epsilon_{str\_LTD}$ | 0.1 | Cortico-striatal learning rate for LTD |
| $\epsilon_{str\_LTP}$ | 0.1 | Cortico-striatal learning rate for LTP |
| $k_{hebb}$ | 0.01 | Constant for hebbian learning term |
| $\epsilon_{cor}$ | 0.00001 | Cortico-cortical learning rate |

## Notes

### Competing Interest Statement

The authors have declared no competing interest.

### Summary of Updates

Abstract length reduction to 250 words

